# Eye lens organoids going simple: characterization of a new 3-dimensional organoid model for lens development and pathology

**DOI:** 10.1101/2023.07.12.548679

**Authors:** Matthieu Duot, Roselyne Viel, Justine Viet, Catherine Le Goff-Gaillard, Luc Paillard, Salil A. Lachke, Carole Gautier-Courteille, David Reboutier

**Author notes:** These authors contributed equally to this work (Matthieu Duot, Roselyne Viel). SAL, CGC and DR are joint senior authors on this work. Corresponding authors (CGC); (DR).

## Abstract

The ocular lens, along with the cornea, focuses light on the retina to generate sharp images. Opacification of the lens, or cataract, is the leading cause of blindness worldwide. Presently, the best approach for cataract treatment is to surgically remove the diseased lens and replace it with an artificial implant. Although effective, this is costly and can have post-surgical complications. Toward identifying alternate treatments, it is imperative to develop organoid models relevant for lens studies and anti-cataract drug screening. Here, we demonstrate that by culturing mouse lens epithelial cells under defined 3-dimensional (3D) culture conditions, it is possible to generate organoids that display optical properties and recapitulate many aspects of lens organization at the tissue, cellular and transcriptomic levels. These 3D cultured lens organoids can be rapidly produced in large amounts. High-throughput RNA-sequencing (RNA-seq) on specific organoid regions isolated by laser capture microdissection (LCM) and immunofluorescence assays demonstrate that these lens organoids display spatiotemporal expression of key lens genes, *e.g.*, *Jag1*, *Pax6*, *Prox1*, *Hsf4* and *Cryab*. Further, these lens organoids are amenable to induction of opacities. Finally, knockdown of a cataract-linked RNA-binding protein encoding gene, *Celf1*, induces opacities in these organoids, indicating their use in rapidly screening for genes functionally relevant to lens biology and cataract. In sum, this lens organoid model represents a compelling new tool to advance the understanding of lens biology and pathology, and can find future use in the rapid screening of compounds aimed at preventing and/or treating cataract.

## Introduction

The lens, in conjunction with the cornea, is responsible for the focusing of light onto the retina, thus creating a clear image (Cvekl and Zhang, 2017). It is a fully transparent biological tissue that involves extreme cell differentiation processes. At the histological level, the lens is composed of an anterior monolayered epithelium containing proliferating cells in the equatorial region that later exit the cell cycle and progressively differentiate into fiber cells (Bassnett and Šikić, 2017). These latter cells form the lens cortex, once the differentiation process is complete. To achieve lens transparency, fiber cells lengthen extensively (∼1000X), produce large amounts of refractive proteins called crystallins, and eliminate their organelles, including their nuclei (Bassnett et al., 2011; Liu et al., 2022). Lens clouding or cataract is the leading cause of blindness worldwide. While the primary reason for the development of cataracts is aging, they can also be induced by environmental factors or have a congenital origin, often triggered by genetic predispositions or abnormalities (Shiels and Hejtmancik, 2021, 2019). To date, the only treatment for cataracts is surgery, which consists of replacing the clouded lens with an artificial implant. Although it is effective, surgery is costly and can have side effects that are far from harmless (Liu et al., 2017). Therefore, efforts to develop drugs to treat cataracts have been initiated (Daszynski et al., 2019; Makley et al., 2015; Zhao et al., 2015). Animal models such as zebrafish (Goishi et al., 2006), Xenopus (Viet et al., 2019), chicken or mammals, namely, rodents, dogs or macaques (Chang, 2016) are used for the study of lens pathophysiology. However, a major bottleneck toward developing anti-cataract drugs remains the lack of an adequate biological model for intensive drug screening.

In recent years, biology and medicine have undergone a revolution with the advent of particular 3-dimensional (3D) cultures called organoids (Zhao et al., 2022). These are *in vitro* cellular models that mimic several aspects of the structure and function of the corresponding organ. Lens epithelium explants were a first generation of 3D lens cultures (O’Connor and McAvoy, 2007). Later on, lentoid bodies, which are 3D cellular structures emerging from various types of 2D stem cells cultures (Anchan et al., 2014; Fu et al., 2017; Yang et al., 2010), or individual micro-lenses grown from lens epithelial cells or pluripotent stem cells (Murphy et al., 2018; Plüss and Kustermann, 2019) were described. Although these models have very interesting molecular and/or optical characteristics, they do not exhibit any particular organization reminiscent of the histology of the lens (Cvekl and Camerino, 2022). Moreover, they often require sequential treatments by individual or combined growth factors and remain tricky and time-consuming to implement. Consequently, they generally do not allow for high-throughput studies.

The goal of the present study was to develop a mammalian organoid lens model that could be generated rapidly and is more convenient to use. As a starting point, we considered a previous paper which shows that lens epithelium can regenerate a functional lens after its ablation in several vertebrate models (Lin et al., 2016). This capacity relies on the presence of lens stem or progenitor cells that sustain self-renewal. Characterization of these cells demonstrated that they express *Pax6* and *Bmi1* and that they are able to spontaneously generate lentoid bodies. The 21EM15 mouse lens <u>e</u>pithelial <u>c</u>ell (LEC) line expresses *Pax6* and *Bmi1* and can spontaneously forms lentoid bodies *in vitro* (Terrell et al., 2015; Weatherbee et al., 2019). However, these lentoid bodies have not been characterized and the culture conditions to controllably induce 21EM15 cells to become such 3D structures have not been defined.

Therefore, in the present study, we sought to derive the culture conditions that could generate lens organoids from 21EM15 cells *en masse*. Further, we sought to undertake their detailed characterization to evaluate their utility in studying genes and pathways relevant to lens biology and pathology. Our work indicates that using simple 3D culture conditions, we can generate numerous lens organoids in a short period of time. These organoids show very interesting optical properties and recapitulate lens physiology at the molecular, histological and cellular levels. In addition, our results demonstrate the possibility to induce various types of opacities, thus mimicking cataract, in these organoids. As a whole, the 21EM15 organoids should therefore provide the lens community with a compelling new model to advance the understanding of lens biology and pathology. From a clinical point of view, these organoids can be potentially used for screening compounds that could have an effect on both the prevention and/or treatment of cataract.

## Results

### 21EM15 spheroids are transparent and have the ability to focus light

To test their ability to grow under 3D culture condition, we seeded 21EM15 cells in 96-well culture plate wells coated with polyhema. Twenty-four hours after seeding, the vast majority of the cells were assembled into round spheroids with no isolated cell being observed. Upon subsequent culture, these spheroids grew in size (Supplemental Figure 1A), acquiring an ovoid asymmetric shape between days 3 and 7. Thereafter, culturing the spheroids beyond day 10 only resulted in limited changes in their overall appearance (Supplemental Figure 1A). At day 10, 21EM15 spheroids are transparent contrary to the spheroids generated with two other epithelial cell lines grown in the same conditions, namely, human epithelial keratinocytes (HaCaT cells) or head and neck squamous cell carcinoma A253 cells (Figure 1A). We then tested the capacity of the 21EM15 spheroids to focus light following a previously described approach (Murphy et al., 2018). Briefly, we imaged the spheroids at different *z*-positions starting from the focus (Figure 1B). A very bright light spot is observed at the center of the 21EM15 spheroids at a specific *z*-position (Figure 1C), suggesting that they had acquired properties to focus light. We quantified the light focusing ability as the ratio of the maximum light intensity at the center of the spheroid to the mean intensity around the spheroid (Figures 1C and D). This ratio reaches values well above 1 at *z*-positions below the focus in 21EM15 spheroids, confirming their capacity to focus light (Figure 1D), not observed in HaCaT spheroids (Supplemental Figure 1B).

**Figure 1.**
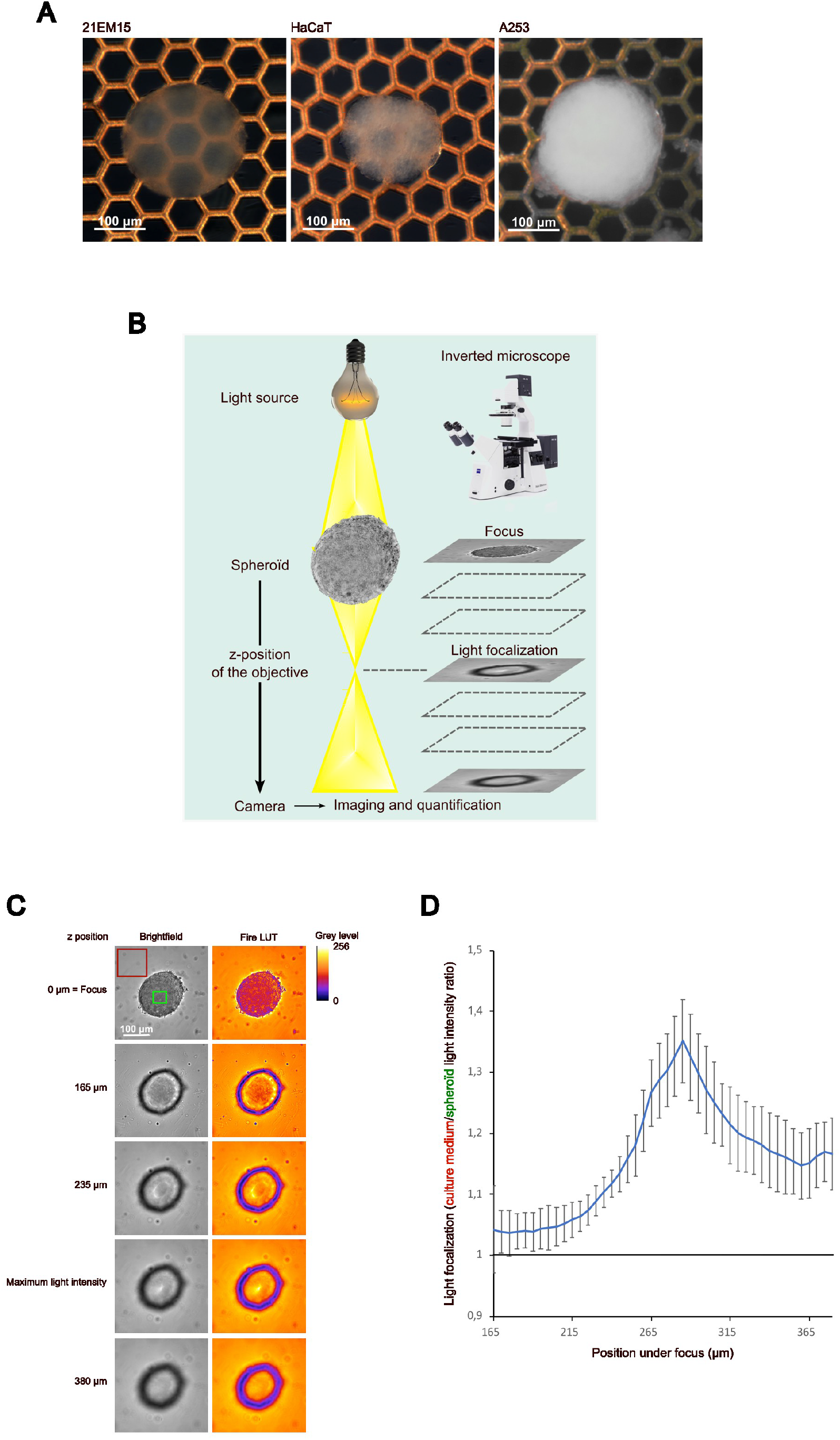
21EM15 spheroids are transparent and can focus light. A, Macroscopic views of 21EM15, HaCaT or A253 10-day old spheroids formed with 10,000 cells. The electron microscopy grid allows evaluation of transparency. B, Schematic of the imaging setup for quantifying the light focusing ability of spheroids. C, Microscopic images of a 21EM15 spheroid showing its ability to transmit and focus light. The mean intensity of light transmitted by the medium and the maximum intensity of light transmitted by the spheroid used for quantification are respectively indicated by the red and the green squares. D, Graph showing the light focusing ability of the 21EM15 spheroids calculated as the ratio between the mean intensity measured in the red square and the maximum intensity measured in the green square. This graph is representative of 5 independent experiments with n=12 spheroids for each experiment. Error bars represent standard deviations. All spheroids were generated from 10,000 cells.

### Transcriptome analysis of 21EM15 spheroids reveals strong similarities with lens development

During their growth, the 21EM15 spheroids acquire asymmetric shape and optical properties to focus light. To gain insights into the molecular modifications associated with these morphological changes and assess whether they may be relevant to lens development, we compared the transcriptome landscape of 21EM15 spheroids with those of mouse lenses at several stages of development. We profiled gene expression in 21EM15 spheroids by 3’ end RNA-sequencing (Supplemental table 1) and we retrieved gene expression levels of mouse lenses from the iSyTE 2.0 database (Kakrana et al., 2018). In iSyTE 2.0, “lens-enrichment” is estimated as the log-ratio of gene expression in the lens to gene expression in the whole embryonic body (WB). Using WB comparative analysis, we similarly estimated gene enrichment in 3D 21EM15 cultures. We compared the 10% most enriched genes in 3D cultures (N = 1032, 10,320 genes in total) with the 10% most enriched genes in E14.5 lenses (N = 1032). We found that 198 genes are present at the overlap of the two datasets, which is far above what was expected by chance (p = 8.7×10^-22^, hypergeometric test). Irrespective of the threshold set to classify the genes as top-enriched (between 0 and 10%), the number of genes observed in the overlap of top-enriched genes in 3D cultures and top-enriched genes in E14.5 lenses largely exceeds that expected (Figure 2A). Nine genes are present in the overlap of top-1% most enriched genes in E14.5 lenses and the top-1% most enriched genes in 3D cultures, whereas only one was expected by chance. Among these genes are the *Cryab*, *Six3*, *Adamtsl4*, *Cp* (encoding ceruloplasmin), *Crim1*, *Dkk3* and *Nupr1*, all of which are known to be directly linked to lens pathophysiology (Figure 2A). We obtained similar results for all lens development stages present in iSyTE 2.0 (E10.5, E12.5, E14.5, E16.5, E17.5, E19.5, data not shown). Together, these results reveal an overlap in expression of key genes between 21EM15 spheroids and normal lenses.

**Figure 2.**
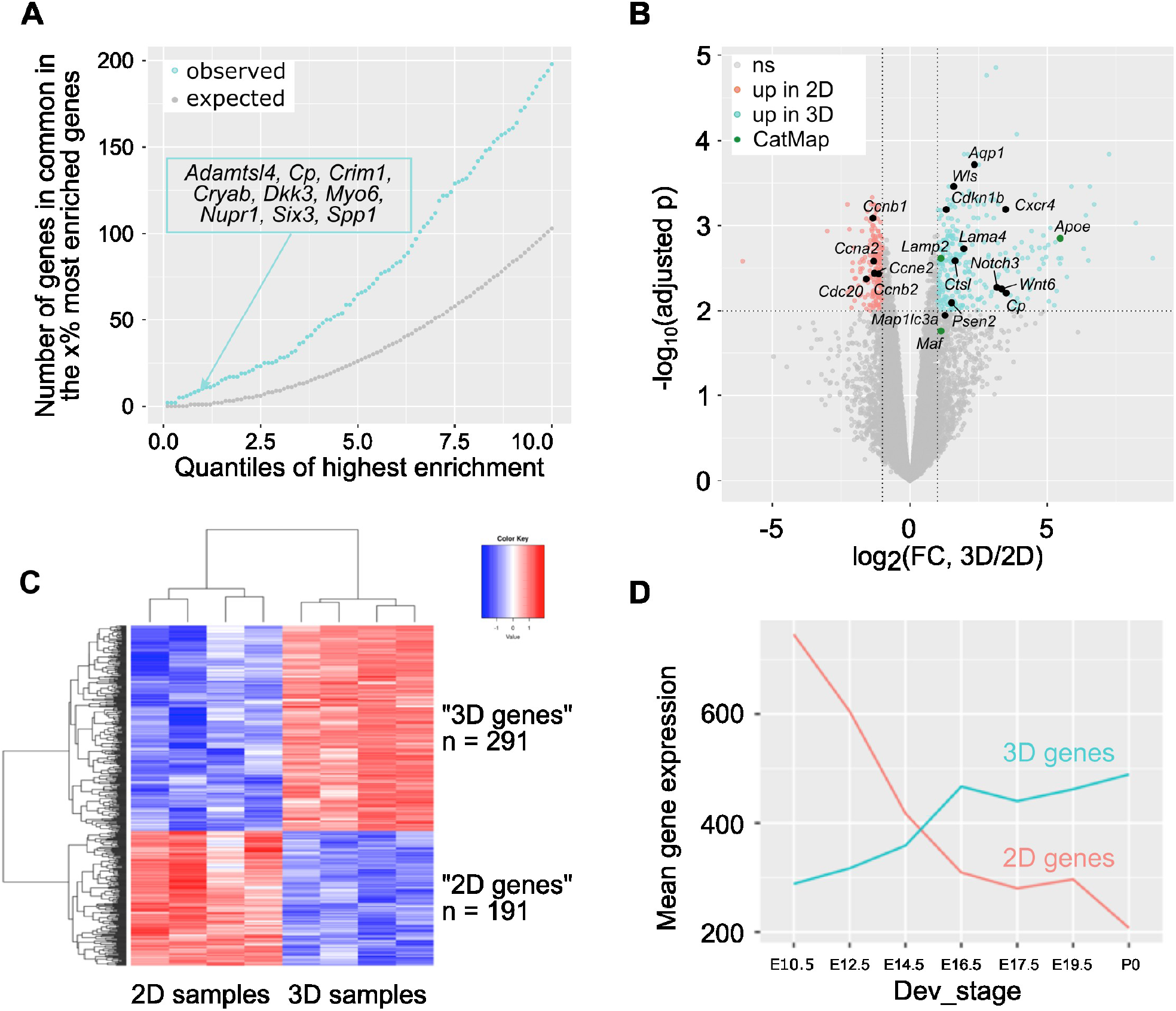
Growing 21EM15 cells in 3D culture conditions captures aspects of gene expression in lens development. A, We ranked all the genes expressed in mouse E14.5 lens and in 21EM15 3D cultures (N = 10,320) based on lens-enrichment (expression in the lens compared to whole body) as described in (Kakrana et al., 2018). We separately listed the percent (x%) of the most enriched genes in the lens and in 3D cultures for several x values ranging from 0 to 10 (X-axis). For each x value, we retrieved the genes in common between the x% most enriched genes in lens and the x% most enriched genes in 3D cultures. The Y-axis shows the number of shared genes for each x value. Blue, the observed number. Grey, the number expected if enrichment in 3D 21EM15 cultures is independent from enrichment in E14.5 lenses. The inset is the list of genes that are in both the 1% most enriched genes in 3D 21EM15 cultures and in the 1% most enriched genes in E14.5 lenses. B, Volcano plot showing the statistical significance (Benjamini-Hochberg adjusted p-value, -log10 scale) against the fold change (log2 scale) of the expression in 2D and 3D samples. This analysis identifies 482 differentially expressed genes (FDR < 0.01 and log2(FC) > 1 in absolute value), among which 291 are upregulated in 3D samples and 191 are upregulated in 2D samples. Genes that are found to be linked to cataract in the database Cat-Map are indicated. C, Heat map of differentially expressed genes in 21EM15 spheroids (3D samples) and 21EM15 2D cultures. D, Mean expression of the 191 “2D genes” and the 291 “3D genes” throughout lens development. In A and D, data for gene expression in lens are from iSyTE 2.0 (Kakrana et al., 2018).

We next wanted to assess the contributions of the culture conditions (3D vs. 2D) on gene expression, especially as it relates to the lens. To do so, we profiled gene expression in 21EM15 2D cultures by 3’ end RNA-sequencing in the same conditions as 3D spheroids (Supplemental Table 1). Principal component analysis and hierarchical clustering analysis showed that the 2D and 3D samples cluster separately from each other (Supplemental Figures 2A and B). We therefore used these datasets to identify differentially expressed genes (DEGs). At FDR = 0.01 and log2(Fold Change) > 1 in absolute value, this analysis uncovered 291 genes that are elevated and 191 genes that are reduced in 3D spheroids (Figure 2B). The elevated 291 genes are referred to as “3D genes” and the reduced 191 genes as “2D genes”. As expected, the heat map of these 482 DEGs shows a clear separation between these two sets of genes (Figure 2C). Several genes elevated under 2D conditions of growth relate to the cell cycle, such as *Ccnb1*, *Ccnb2*, *Ccnb1*, *Ccne2*, *Cdc20* (Figure 2B). This indicates that culturing 21EM15 in 3D conditions reduces the expression of cell cycle genes comparted to growth under 2D conditions. Conversely, several genes elevated under 3D growth conditions are relevant to lens development (Figure 2B). These include *Ank2*, *Aqp1*, *Atp1b1*, *Cap2*, *Eya1*, *Fundc1*, *Fzd6*, *Hsf4*, *Jag1*, *Maf*, *Meis1*, *Prox1*, *Tdrd7* and *Wls*.

Finally, we used iSyTE 2.0 to examine the expression of these DEGs in normal lens development (Kakrana et al., 2018). On average, 2D genes are more expressed than 3D genes in early lens developmental stages (*e.g.* E10.5, when the lens placode has invaginated into a “lens pit”), and their expression decreases as the lens progresses in development (Figure 2D). Conversely, the mean expression of 3D genes increases progressively in normal lens development (Figure 2D). Hence, switching the culture conditions of 21EM15 cells from 2D to 3D spheroids partly recapitulates gene expression changes in normal lens development. Together, these data show that culturing 21EM15 in 3D conditions reinforces their similarity with normal mouse lenses, and in particular at later stages of development.

### 21EM15 3D cultured cells form multilayered lens organoids

The lens is surrounded by a basal lamina called the capsule. Under the capsule and starting from the anterior pole of the lens, there is a quiescent epithelium, containing cells that proliferate in the “germinal zone” and exit the cell cycle at the “transition zone” to initiate differentiation into fiber cells that contribute to the bulk of the lens. Differentiation into fibers cells is characterized by the remodeling of the cytoskeleton leading to elongation of the cells, accompanied by high levels of expression of key lens proteins (e.g., crystallins, membrane proteins, etc.) and the progressive loss of cellular organelles (Bassnett et al., 2011; Liu et al., 2022). As 21EM15 cells cultured as 3D spheroids express genes associated with lens differentiation, we wanted to characterize their structure to identify whether different cell types emerge through a process of differentiation. Histological analysis shows that twelve hours after seeding under 3D conditions, 21EM15 cells cluster to form a non-organized flattened structure (Figure 3A). Twenty-four hours after seeding, the structure is spherical with a rather homogenous cellular content. Ten days after seeding, the spheroids show a different appearance, with distinct zones: the external zone with round nuclei, the intermediate zone with elongated nuclei, and the internal zone with cells characterized by an intense pink cytoplasm and small compacted nuclei (Figure 3A). Importantly, the internal zone is off-centered, revealing that the initial central symmetry of the spheroid was broken during the 10-day culture (Figure 3A). This is in line with previous observations that the spheroid acquires an ovoid shape after a few days of culture (Supplemental Figure 1A).

**Figure 3.**
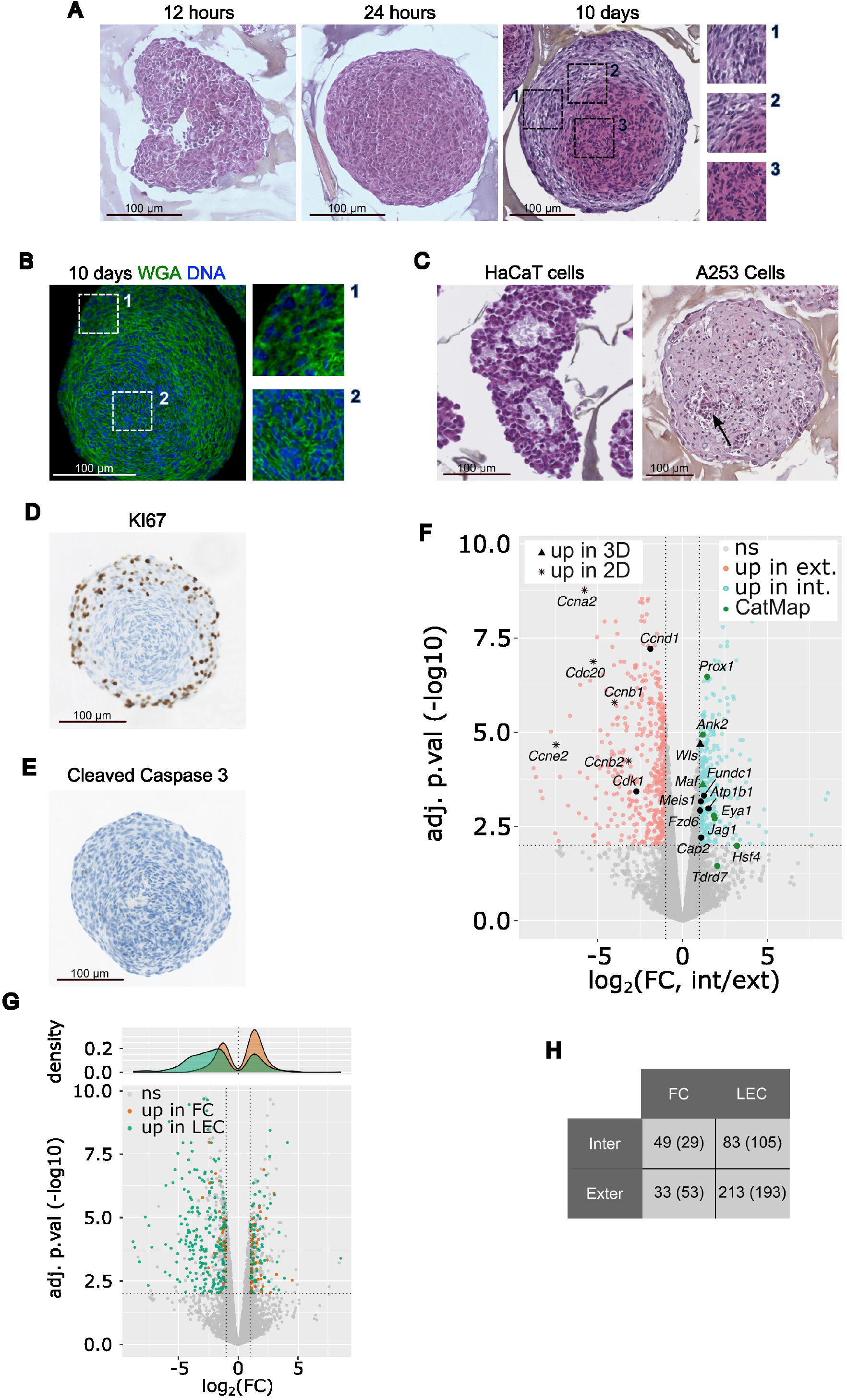
21EM15 LECs 3D cultures differentiate to form multilayered lens organoids. A, Histological analysis of 21EM15 spheroids grown for 12 hours, 24 hours or 10 days, stained with Hematoxylin, Eosin and Safran (HES). The insets show the three different histological regions. B, Microscopic image of a 21EM15 spheroid stained with Wheat Germ Agglutinin (WGA) to show distinct cellular boundaries. Insets show details of the outer (1) or inner regions (2). C, Histological analysis of HaCaT or A253 spheroids grown for 10 days (HES staining). The arrow shows a necrotic region. D, Histological analysis showing the localization of KI67 in a 10-day old 21EM15 spheroid. E, Cleaved Caspase 3 staining of 10-day old 21EM15 spheroid, revealing an absence of cells undergoing apoptosis. For figures A-E, data are representative of at least 3 independent experiments, n=30 organoids. F, Volcano plot showing the statistical significance (Benjamini-Hochberg adjusted p-value, -log10 scale) against the fold change (FC, log2 scale) of the expression in internal and external regions microdissected from 21EM15 spheroids. This analysis identifies 793 differentially expressed genes (FDR < 0.01 and log2(FC) > 1 in absolute values), among which 328 are upregulated in internal regions and 465 are upregulated in external regions. “Up in 2D” and “Up in 3D” correspond to genes respectively up in 2D or up in 3D in figure 2B. Black solid symbols correspond to genes that are of particular significance regarding the lens. G, Lower panel, same volcano plot as in F, with genes that are overexpressed in microdissected lens epithelial cells (LEC) and fiber cells (FC) (Zhao et al., 2018) colored in green and orange, respectively. Higher panel, density plot showing the distribution of fold changes (log2 scale) for FC and LEC genes. H, Contingency table showing the number of genes that are enriched in FC or LEC compared to internal and external regions. The numbers in the brackets represent the value that is expected in the event that the enrichment in FC or LEC was unrelated to the enrichment in internal or external regions, respectively. The difference between the expected and the observed value indicates that a higher number of genes than expected are found to be enriched both in FC and internal regions, as well as in LEC and external regions.

To evaluate cell compartmentalization in the core of the spheroid we visualized cells boundaries, using WGA (wheat germ agglutinin), a membrane marker. The core of the spheroid is composed of highly compacted cells when compared to the cortex (Figure 3B). This indicates that the cells located at the central region of the spheroid undergo a phenomenon of packing. As controls, histological sections of HaCaT and A253 spheroids were generated (Figure 3C). After 10 days of culture, specific organization of cells was not observed in these controls. HaCaT spheroids are made of cells roughly aggregated and cavities likely filled with extracellular matrix, while A253 spheroids are made of homogenously distributed cells with a clear pink cytoplasm and some spots of necrotic cells (Figure 3C). As 21EM15 spheroids constantly grow over a period of more than 10 days (see Supplemental Figure 1B), we wanted to determine which cells were responsible for the growth. Staining with the KI67 antigen, an established marker of proliferation, shows that proliferating cells are essentially localized in the external region of the spheroid and that cells stop proliferating once their nuclei are elongated (Figure 3D). The observation that only a few cells are able to proliferate is consistent with the finding that many cell cycle-related genes are down-regulated in 3D cultures (Figure 2B). No cell undergoing apoptosis was observed after 10 days of culture (Figure 3E).

The above data indicate that 21EM15 spheroids are made up of at least 3 distinct regions. We next sought to determine the gene expression landscape within the different layers of the spheroids. Toward this goal, the most internal and external regions were isolated by laser capture microdissection (LCM) and subjected to 3’ end RNA-sequencing (Supplemental table 2). Due to technical limitations, we were unable to microdissect the intermediate region. We retained *n*=3 external region samples and *n*=4 internal region samples based on PCA (Supplemental Figure 3A). We identified 793 DEG (FDR = 0.01, and log2(Fold Change) > 1 in absolute value). Of these, 465 exhibit enriched expression in the external region and 328 exhibit enriched expression in the internal region (Figure 3F). The heat map of these 793 DEGs separates “external genes” from “internal genes” (Supplemental Figure 3B). Among the genes that are overexpressed in the internal region, we found genes known to be expressed in fiber cells like *Ank2*, *Atp1b1*, *Cap2*, *Eya1*, *Fundc1*, *Fzd6*, *Hsf4*, *Jag1*, *Maf*, *Meis1*, *Prox1*, *Tdrd7*, *Wls*. Among the genes that are overexpressed in the external region, we found genes known to be expressed in lens epithelial cells like *Ccna2*, *Ccnb1*, *Ccnb2*, *Ccnd1*, *Ccne2*, *Cdc20*, *Cdk1*.

To globally assess the resemblance of the internal and external regions of 21EM15 spheroids with lens fiber and epithelial cells, respectively, we retrieved the transcriptomic data from microdissected E14.5 epithelial cells and lens fiber cells (Zhao et al., 2018). Of the 793 DEG, 378 are also enriched either in E14.5 lens epithelium or fiber cells. The “external genes” (negative log_2_(FC) in Figure 3G), are enriched in lens epithelial genes (green spots), whereas “internal genes” (positive log_2_(FC) in Figure 3G) are enriched in lens fiber cells (orange spots). The contingency table shown in Figure 3H confirm this bias (p = 2.1*10^-6^, chi-square test). These data confirm that the transcriptome of the internal region resembles that of lens fiber cells and the transcriptome of the external region that of lens epithelial cells. Further, while 2D cultures of 21EM15 cells have overlapping expression with lens epithelial cells, growing the cells in 3D commits the internal cells toward a differentiation program overlapping with lens fiber cells. Taken together, our results show that the 3D 21EM15 cultures self-organize and establish different cell types expressing specific gene sets, thus mimicking certain aspects of lens development. Moreover, these structures are able to break their original central symmetry to establish axial symmetry, which is characteristic of organoid development (Anand et al., 2023; Brassard and Lutolf, 2019; Serra et al., 2019). Thus, henceforth, the 3D 21EM15 cultures will be referred to as 21EM15 “organoids”.

### Morphological organization of 21EM15 organoids partially recapitulates lens patterning

Different regions or cell types within the lens can be characterized by expression of specific markers. From the transcriptomic studies described above, we identified a subset of key genes involved in lens development to be differentially expressed between the inner and outer regions (see Supplemental Table 2). Our next objective was to confirm that key lens genes have specific expression patterns relevant to the organization of a whole lens. For this purpose, we examined the expression and the localization of structural components such as Laminin, a major component of basal lamina including the lens capsule, and αB-Crystallin (CRYAB), a major component of lens fiber cells in later developmental stages (Cvekl et al., 2015; DeDreu et al., 2021). We also examined transcription factors such as PAX6, PROX1 (elevated in fiber cells) and signaling molecules as JAG1 (Cvekl and Zhang, 2017; Duncan et al., 2002; Garg et al., 2020). Laminin was found to be present in 2 or 3 layers of the outermost cells (Figure 4A). It is also present in the most peripheral part of the lens, but only in the form of a single outer basal lamina (DeDreu et al., 2021). Immunostaining showed that nuclear PAX6 is present in an internal region surrounding the central core of the organoid composed of highly compacted cells (Figure 4B). Consistent with symmetry breaking, it then extends toward the central axis. JAG1 is also found in an asymmetric distribution, as it is enriched in the membranes of cells localized in two lateral areas surrounding the central axis of the organoid (Figure 4B). PROX1 is present in two regions: in the cytoplasm of cells that most strongly express *Jag1*, and in the nuclei of cells that more weakly express *Jag1* and are localized along the central axis (Figure 4B and C). Finally, αB-Crystallin is low/absent in the external cell layers but high in the cortex and the core of the organoid with both cytoplasmic and nuclear localization (Figure 4D).

**Figure 4.**
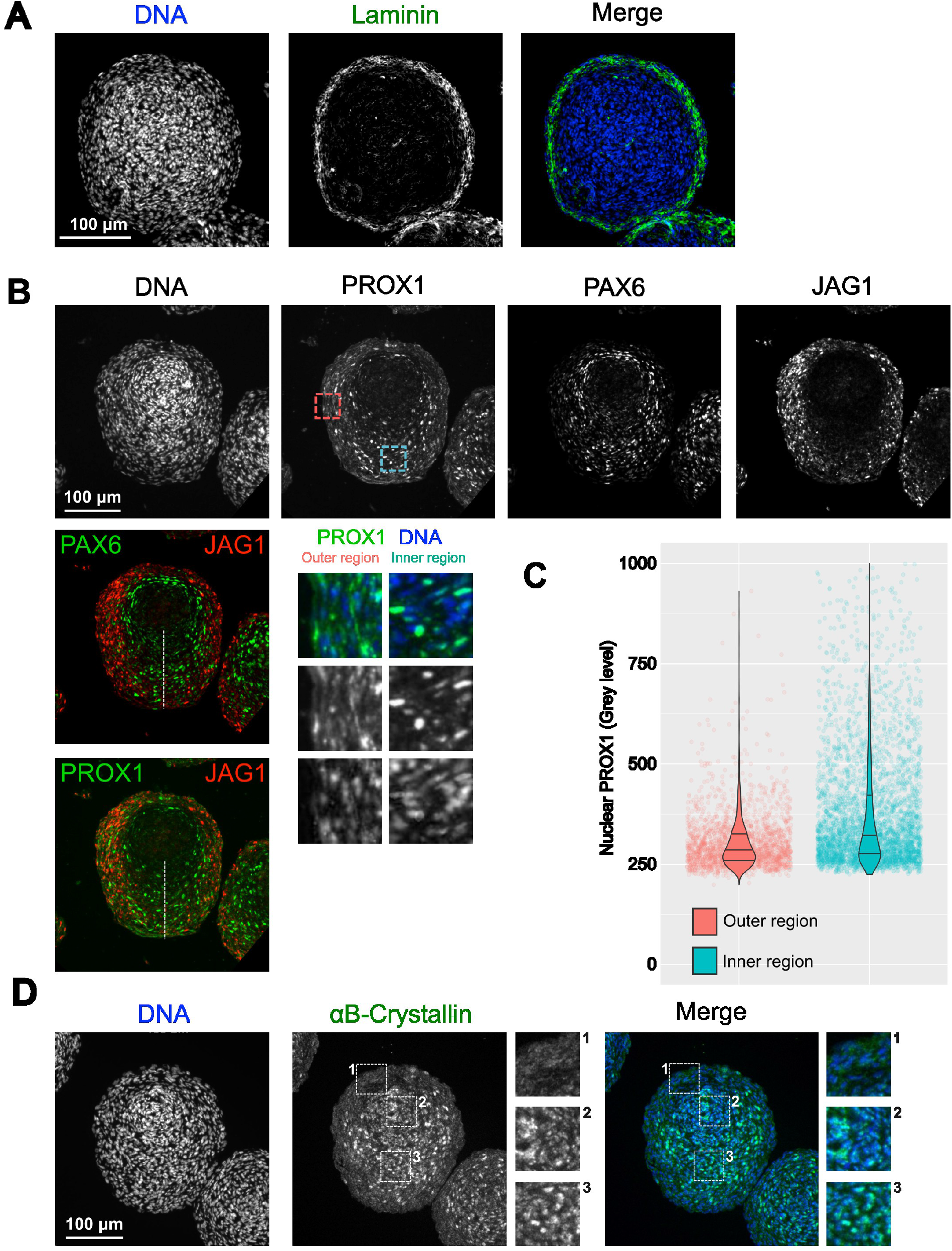
Morphological organization of 21EM15 organoids partially recapitulates eye lens patterning. A, Immunofluorescence (IF) microscopy reveals the localization of Laminin on the outer region of the organoid. B, Multiplex microscopic images indicates the localization of the lens expressed proteins PAX6, PROX1 and JAG1. The red and blue squares correspond to the typical areas (respectively outer and inner regions) used to display the magnification insets presented below the grey level image of PROX1 and to quantify the PROX1 nuclear labeling shown in C. The white dashed line symbolizes the central axis of the organoid. Violin plots combined with jittered scattered plots showing that PROX1 is enriched in nuclei of cells located in the inner region. This graph is representative of three independent experiments, n=30 organoids. *p* < 2.2 x 10^-16^, Wilcoxon rank sum test with continuity correction. D, IF images showing the localization of αB-Crystallin. Insets show the enlargement of the outer region, the central axis and the core of the organoid. Data are representative of at least three independent experiments, n=30 organoids.

One of the most striking characteristics of fiber cell differentiation is the progressive degradation of cellular organelles such as nuclei and mitochondria (Brennan et al., 2021). In the lens, these various events can be highlighted by the observation of components of the nuclear envelope, of the mitochondria or by the expression of specific transcription factors (Costello et al., 2013; Cui et al., 2016; Imai et al., 2010; Morishita et al., 2013). We sought to examine whether similar cellular changes occurred in 21EM15 organoids. We find that Lamin-B1 (LMNB1), a component of the nuclear envelope, progressively disappears from the exterior to the interior of the organoid (Figure 5A). While Lamin-B1 labeling surrounds the nucleus in a continuous manner in the outermost cells, this labeling becomes more and more discontinuous toward the inner region of the organoid, until it completely disappears. Concomitantly, we observed a gradual change of nuclei shape accompanied by chromatin compaction evoking pyknosis (Figure 5B). These nuclei are not transcriptionally active in cells located in the center of the organoids (Figure 5C) as indicated by the progressive loss of nuclear Fibrillarin (FBL), which is considered as a marker of transcriptional status of fiber cell nuclei (Sandilands et al., 2002). Mitochondria are also in a process of degradation as indicated by the decrease in TOMM20 labelling, a constituent of the mitochondria external membrane, toward the center of the organoid (Figure 5D). Conversely, the expression of *Hsf4,* a gene encoding a transcription factor whose downstream targets are considered to be involved in organelle degradation (Cui et al., 2016, 2013), increases in the core region relative to the outer region (Figures 5E and F).

**Figure 5.**
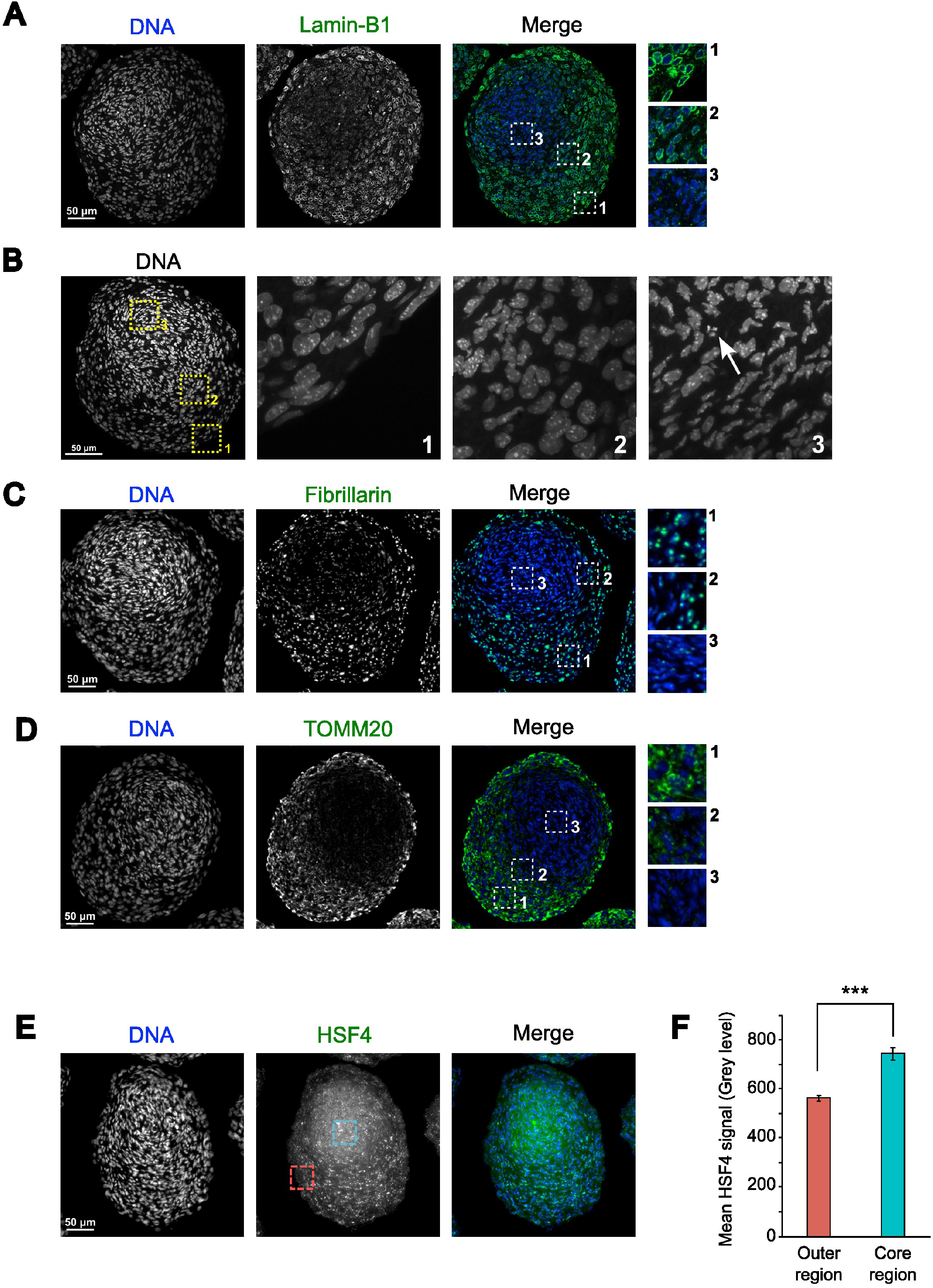
Cells located in the core of lens organoid are engaged in a process of organelle degradation. A, IF microscopy images of 10-day old lens organoid showing the localization of Lamin-B1. B, Microscopy images showing the appearance of nuclei in three different areas of an organoid. The arrow points out a nucleus with pyknotic features. C, Nuclei located in the core of the organoid show reduced transcriptional activity, as evidenced by decreased levels of Fibrillarin, which has previously been used as a marker of the transcriptional state of lens fiber cells. D, The decrease in TOMM20 labeling indicates that the organoid core cells are engaged in mitochondrial degradation. E, IF microscopy images showing the localization of HSF4. The red and blue squares correspond to the typical areas (respectively outer and inner regions) used to quantify the HSF4 mean signal quantified in F. F, Histogram presenting the quantification of the mean HSF4 signal in the outer and the core regions of the organoids. This graph is representative of three independent experiments, n=30 organoids. Asterisks indicate a *p*-value < 0.001. Error bars represent standard deviations.

Taken together, these results suggest that similar to the cellular and morphological changes accompanying lens development, the 21EM15 organoids are organized into specific expression domains for key lens proteins like αB-Crystallin, PAX6, PROX1 and JAG1. Moreover, the cells lying in the internal-most region commit to a process of organelle degradation reminiscent of what is observed in the whole lens. Our results thus show that 21EM15 organoids recapitulates specific molecular aspects and morphological organization of the lens.

### 21EM15 organoids model lens cataract

As the organoids described above present interesting optical, morphological, histological, molecular and functional characteristics, we explored if they could be utilized as a model to uncover the pathophysiology of the lens. Cataract can be induced by H_2_O_2_ or hypertonic NaCl treatments in dissected lens (Ruiss et al., 2021). Therefore, we incubated 8-day old 21EM15 organoids for 48 hours with these compounds (at previously used concentrations) and evaluated their transparency and light focusing ability. We found that H_2_O_2_ does not trigger changes in transparency or light focusing ability for concentrations ranging from 0 to 350 μM, but organoids become opaque and cease to transmit light at concentrations above 500 μM (Figures 6A and B). Increasing concentrations of NaCl (from 1.25% to 1.7%) gradually reduce organoid transparency (Figure 6A), but only the highest concentration has a significant impact on light focusing ability (Figure 6C). We next sought to examine whether the 21EM15 organoid model could be applied for testing function of genes associated with cataract. We previously showed that *Celf1* deletion in a germline or lens conditional manner causes early-onset cataract in mice (Siddam et al., 2018). We therefore tested the transparency and light focusing ability of organoids made from 21EM15 cells stably expressing a shRNA targeting the *Celf1* gene (Aryal et al., 2020; Siddam et al., 2018). Interestingly, *Celf1* knockdown reduces organoids transparency, as observed in mice deficient for *Celf1* (Figure 6D). It also reduces the light focusing property of 21EM15 organoids (Figure 6E). All together, these results demonstrate that 21EM15 organoids are a new model that can be used to study cataract.

**Figure 6.**
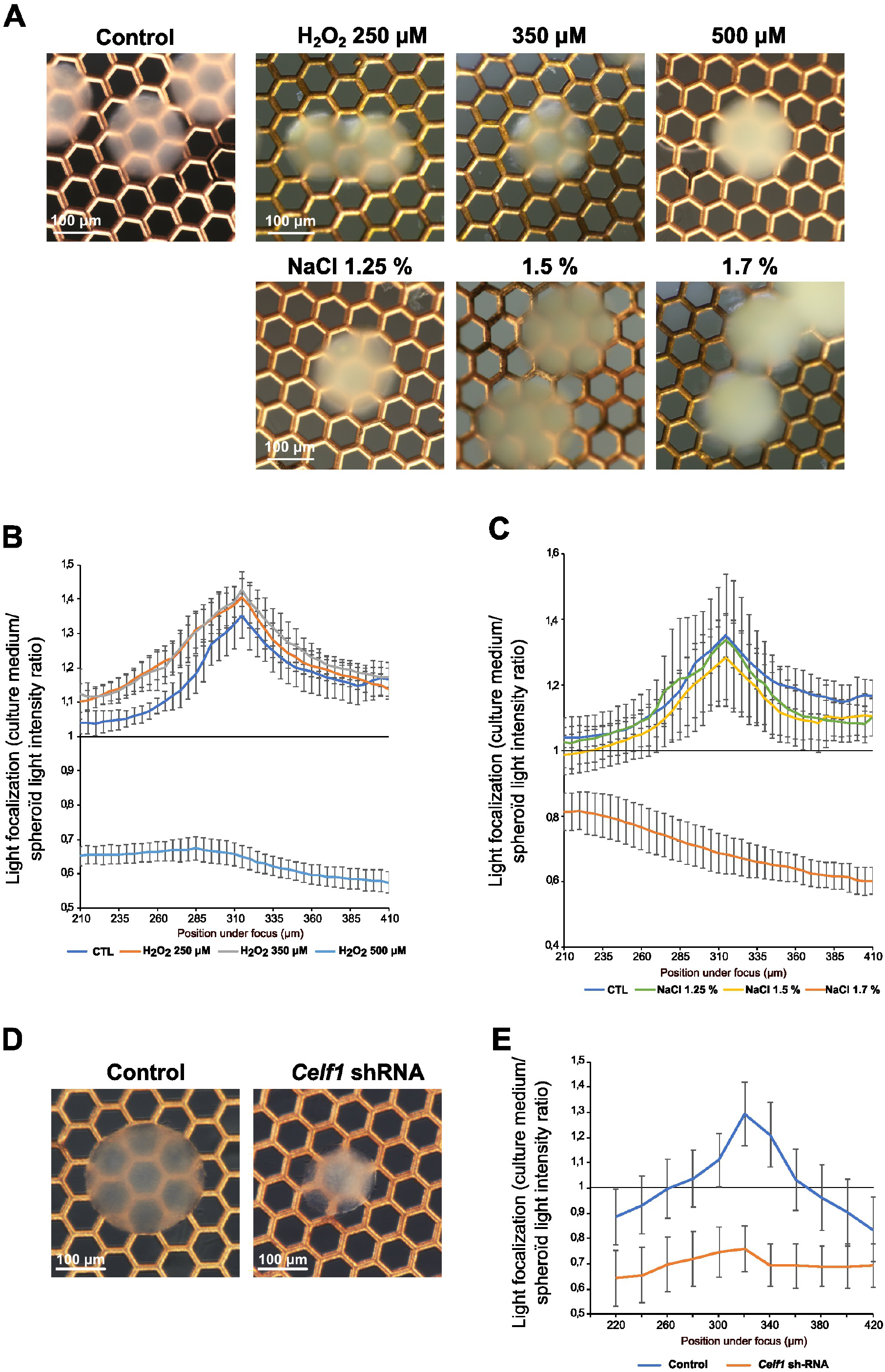
21EM15 lens organoids can be induced to develop opacity. A, Macroscopic views of 10-day old lens organoids treated with the indicated concentrations of H_2_O_2_ or NaCl. The electron microscopy grid allows evaluation of transparency. B, Graph showing the light-focusing ability of the 21EM15 lens organoids treated with increasing concentrations of H_2_O_2_. C, Graph showing the light-focusing ability of the 21EM15 lens organoids treated with increasing concentrations of NaCl. D, Macroscopic views of lens organoids expressing control or *Celf1*-targeting shRNA. The electron microscopy grid allows evaluation of transparency. E, Graph showing the light-focusing ability of the 21EM15 lens organoids expressing control or *Celf1*-targetting shRNA. Graphs B, C and E are representative of three independent experiments with n=12 organoids 10-day old for each experiment. Error bars represent standard deviations.

## Discussion

In the present study, we have developed a mouse 3D lens model that can be rapidly generated and can be applied for studying processes relevant to lens biology. It has specific optical properties, including transparency and light-focusing ability (Figure 1). We characterized this model by a combination of histological, transcriptomic and immunohistochemical/immunofluorescence approaches. This analysis revealed that, similarly to the lens, 21EM15 lens organoids comprises three main regions, a peripheral layer, an intermediate part and a core region.

When grown in 2D culture conditions, 21EM15 cells express typical LEC genes (Terrell et al., 2015). In the present study, we used 3’end RNA-seq to profile the transcriptomes of 21EM15 2D and 3D cultures, and of laser-micro-dissected internal and external regions of the organoids. These analyses show that a subset of genes associated with lens development and fiber cell differentiation are induced under 3D culture conditions (Figure 2D). Among this set of genes we found *Six3*, a well characterized lens development gene (Oliver et al., 1996), *Cp* that is typically expressed in lens epithelium (Harned et al., 2006), *Cryab*, *Crim1* and *Nupr1* that are expressed in lens fiber cells (Anand et al., 2018; Cvekl et al., 2015; Kakrana et al., 2018; Tam et al., 2018) and *Dkk3* that is a component of the Wnt signaling pathway involved in lens development (Wang et al., 2018) (Figure 2B). RNA-seq analysis also shows that the genes elevated in the external region of the lens organoids are enriched for candidates that are related to the cell cycle function (*Ccna2*, *Ccnb1*, *Ccnb2*, *Ccnd1*, *Ccne2*, *Cdc20*, *Cdk1*), whereas those elevated in the internal region are enriched for candidates involved in the Pax6 regulatory pathway (*Cap2*, *Meis1*), Notch and Wnt signaling (*Jag1*, *Wls*, *Fzd6*), Maf pathway (*Maf*, *Eya1*), lens fiber cell morphology and physiology (*Ank2*, *Atp1b1*, *Prox1*, *Tdrd7*) and organelle degradation (*Hsf4*, *Fundc1*) (Figure 3F and Supplemental table 2). These results are relevant to lens organization, as the anterior epithelium is involved in lens growth and the cortex is the place where fiber cells progressively differentiate (Bassnett et al., 2011; Liu et al., 2022).

Accordingly, KI67 staining shows that only the external cells are proliferative (Figure 3D). We also found the outermost region to be positive for Laminin staining (Figure 4A). Laminin is a glycoprotein specific of basal lamina. It is present in the capsule, a cell-free structure around the lens (DeDreu et al., 2021). In 21EM15 organoids, laminin appears to be secreted by cells in the first two or three layers that are encompassed into this extracellular matrix. We were unable to detect any E-cadherin labeling in these external-most cell layers, suggesting that organoid epithelial cells do not fully recapitulate properties of lens anterior epithelium, which express E-cadherin (Pontoriero et al., 2009) (data not shown). This suggests that while 21EM15 cells are able to proliferate or enter the early stages of fiber cell differentiation, they cannot become true epithelial cells in the culture conditions that we used. One possible explanation relates to the fact that 21EM15 cells were likely selected based on their ability to proliferate rapidly (Terrell et al., 2015). Although it is now established that the anterior epithelium contains stem cells (Bassnett and Šikić, 2017; Lin et al., 2016), the 21EM15 cells probably originate from the germinative zone and are therefore engaged in the early stages of fiber cell differentiation process, preventing them from committing to the epithelial fate. Interestingly, they express *Pax6* and *Bmi1*, which are required for lens epithelial cells to regenerate a functional lens (Lin et al., 2016), but also typical stem cell marker genes like *Nes*, *Chrd*, *Sox4, Sox9, Sox12* and *Klf4* (Terrell et al., 2015). They are also able to spontaneously form lentoid bodies (Terrell et al., 2015; Weatherbee et al., 2019). Finally, we show here, based on histological analysis, that symmetry is broken in 3D 21EM15 cell cultures (Figures 3A, 4, 5). Symmetry breaking is a general hallmark of organoids development (Anand et al., 2023; Brassard and Lutolf, 2019; Serra et al., 2019). Taken together, these observations suggest that 21EM15 cells possess stem cell-like properties accounting for their ability to form lens organoids.

A key property of lens fiber cells is the elimination of organelles, as they are potential sources of light scattering. We tested if this also applies to the core region of 21EM15 organoids. Several gene regulatory networks involved in autophagy or nuclear degradation in the lens have been identified (Bassnett, 2009; Brennan et al., 2021). One of them involves the transcription factor HSF4, a major regulator of membrane organelle degradation in the lens (Cui et al., 2016, 2013; Fujimoto et al., 2004; Huang et al., 2015). Accordingly, we found in immunostaining experiments that HSF4 is more abundant in the core than in the peripheral region of the organoid (Figure 5E). At the RNA level, *Bnip3l*, *Lamp1*, *Fundc1*, and *Smurf1*, which are also involved in organelle degradation (Bassnett, 2009; Brennan et al., 2021), also exhibit elevated expression in the core region than in the outer region of organoids (Supplemental Table 2). The high expression of these genes is accompanied by an apparently complete degradation of the mitochondria, as inferred from the loss of TOMM20 staining, a mitochondrial marker commonly used to assess mitophagy in various cell types including lens fiber cells (Chan et al., 2011; Costello et al., 2013; Morishita et al., 2013) (Figure 5D). The shape of the nuclei is also strongly affected in the organoid core, with some nuclei clearly showing a pyknosis-like appearance (Figure 5B). Pyknotic nuclei are characteristic of various types of terminal cell differentiation processes requiring nucleus degradation, as in red cells or lens fiber cells (McAvoy and Richardson, 1986). In addition to organelle degradation, fiber cell differentiation is also featured by a strong expression of crystallins, which involves PAX6 and to some extent, HSF4 (Cvekl et al., 2015). αb-Crystallin (*Cryab*) is poorly expressed in the outer region whereas it is enriched in the central region where *Pax6* is expressed, as revealed by both 3’end RNAseq on laser-microdissected regions (Supplemental Table 2, FDR = 0.03) and IF (Figure 4D). This is similar to the expression pattern of αb-Crystallin in wild-type lenses, wherein it is high in fiber cells compared to epithelial cells in later stages of development (Zhao et al., 2018). Together, these data show that, while the outermost layer of 21EM15 organoids resembles lens epithelial cells, their core region resembles fiber cells.

In the lens, the transition zone is the area where cells of the anterior epithelium exit the cell cycle and begin differentiation into fiber cells that contribute to the bulk of the lens tissue. Beyond the transition zone, cells sequentially elongate and degrade their organelles while expressing key lens development master genes. We were unable to profile the transcriptome of the intermediate region of 21EM15 organoids due to technical limitations in laser microdissection. However, we gained significant insights into this zone by immunostaining of PAX6, JAG1 and PROX1 (Figure 4). These key genes function to orchestrate the development of these different regions of the lens (Cvekl and Zhang, 2017; Mochizuki and Masai, 2014). The function of JAG1 is to keep epithelial cells from the germinative zone undifferentiated and proliferating, by activating Notch signaling (Mochizuki and Masai, 2014; Rowan et al., 2008; Saravanamuthu et al., 2009). PAX6 and PROX1 respectively trigger cell cycle exit and are associated with expression of crystallins and cell elongation during secondary fiber cells differentiation (Audette et al., 2016; Cvekl and Ashery-Padan, 2014; Shaham et al., 2012, 2009; Wigle et al., 1999). PROX1 exhibits dynamic expression and localization in lens cells. PROX1 is first located in the cytoplasm in the anterior epithelium, and then becomes nuclear in the transition zone where it is involved in orchestrating fiber cell elongation and differentiation (Duncan et al., 2002; Wigle et al., 1999). 21EM15 organoids show an interesting distribution of these 3 key genes. JAG1 is consistently localized in the lateral regions and severely reduced near the central axis, in contrast to PAX6, which is highly expressed in a ring-like manner surrounding the organoid core, and also in the central axis. The changes in PROX1 localization seem to recapitulate its endogenous lens expression. While there is diffused cytoplasmic labeling of PROX1 in the *Jag1*-expressing region, PROX1 becomes nuclear along the central axis, a region where *Pax6* is quite highly expressed. It is interesting to note that the cells that exhibit nuclear PROX1 are located in the area where the cells are most elongated, indicating that the organoids recapitulate this functional aspect of PROX1 similar to endogenous lens development (Figure 3A right and Figure 4B).

Overall, these results show that 21EM15 organoids recapitulate several aspects of lens organization, gene expression profiles, biological processes or optical properties. We therefore summarized these data in a model (Figure 7). In this model, we were unable to formally identify any canonical epithelium. We termed the outermost zone comprising the first layers of cells that are proliferative and are embedded in Laminin as the “capsuloid”. The capsuloid likely corresponds to the fusion of the capsule and the germinative zone of the lens epithelium. The lateral region expressing Jag1 and Ki67 is probably a mix of more or less proliferating cells (as indicated by their expression of Ki67) and early differentiating cells (expressing *Jag1*). Although cells from the germinative zone do not express *Jag1*, it is tempting to compare this region of the organoid to the germinative zone in the lens, where cells that express *Jag1* would prevent cell cycle exit and fiber cell differentiation of proliferating cells. In a more internal region, between the germinative zone and the central axis of the organoid, cells begin to express *Pax6*, *Cryab*, and PROX1 becomes nuclear. This region is characteristic of the transition zone in endogenous lens development, where the cells progressively engage in the process of fiber cell differentiation. Interestingly, along the central axis and around the organoid nucleus, the cells no longer express *Jag1*. However, these cells still express *Pax6*, PROX1 is nuclear, and most importantly, exhibit elongation (Figure 4B). Cells in the central-most region exhibit a very intensely stained “pink” cytoplasm in histological analysis, reminiscent of the lens, with pyknotic nuclei and strong expression of *Hsf4* and *Cryab*. They also show features consistent with the degradation of their organelles (loss of Lamin B, TOMM20 and fibrillarin markers). All of these data suggest ongoing cellular and molecular processes in the lens organoids that contribute to transparency.

**Figure 7.**
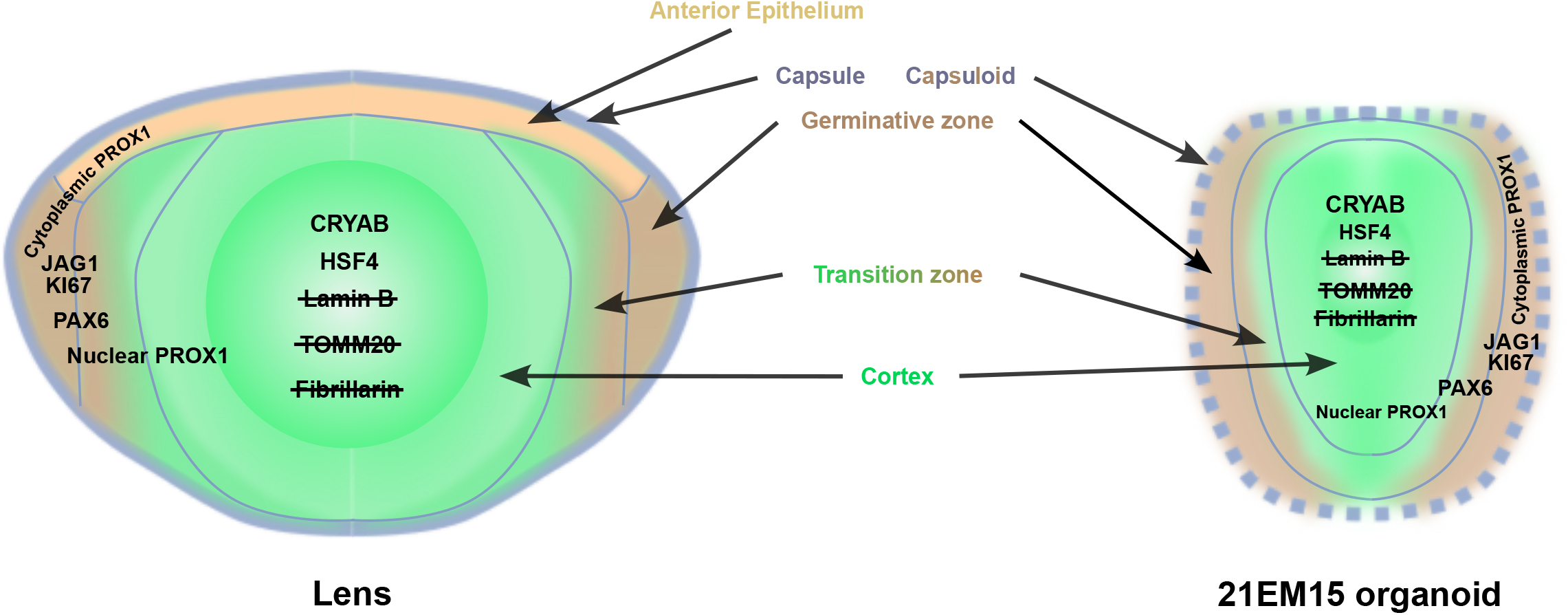
Model for 21EM15 lens organoid organization. This model shows that, unlike the lens, 21EM15 organoids lack the typical lens capsule and anterior epithelium. Instead, there is a zone that we term “capsuloid”, comprising the outer layers of proliferative cells embedded in a laminin layer. Further inwards and from the outside in, 21EM15 organoids recapitulate several aspects of the organization of the lens, with a region corresponding to the germinal zone, where *Ki67* and *Jag1* are expressed, followed by a transition zone that expresses *Pax6* and where PROX1 progressively becomes nuclear. Along the central axis of the organoid, PROX1 is predominantly nuclear and cells progressively degrade their organelles, such as nuclei (loss of Lamin B and Fibrillarin) and mitochondria (loss of TOMM20 signal), and begin to express *Hsf4* and *Cryab*. These events correspond closely to those described in the normal lens cortex.

Finally, we addressed the suitability of 21EM15 organoids as a tool for studying lens opacity or cataracts (Figure 6). We observed that two previously established treatments to induce cataract, namely exposure to H_2_O_2_ and NaCl, also are able to induce opacity in 21EM15 organoids. This is significant as previously, H_2_O_2_ was shown to model the events leading to age-related cataract (Hernebring et al., 2021; Spector, 1995; Spector et al., 1993). Further, treatment with hyperosmotic NaCl was shown to trigger osmotic stress and disrupts fluid balance of lens, frequently associated with dry eye disease and diabetic cataracts (Butler, 1994; Gu et al., 2016; Obrosova et al., 2010). Thus, 21EM15 organoids could find application in further understanding how these processes induce cataract. Finally, we also observed that knockdown of the cataract-linked RNA-binding protein encoding gene, *Celf1*, reduces 21EM15 organoids transparency and light focusing properties. CELF1 is involved in regulation of mRNA splicing, stability or translation (Dasgupta and Ladd, 2012), and its inactivation in mice leads to misregulation of post-transcriptional gene expression control in the lens and cataract (Aryal et al., 2020; Siddam et al., 2023, 2018). These results indicate that 21EM15 organoids respond to cataractogenic conditions representative of a wide range of lens-related etiologies (*e.g.*, age-related cataract, diabetic cataract, genetic cataract) and thus can be used in further advancing knowledge on lens pathology.

In conclusion, our work presents a new mouse organoid model, that is effective and easy to set up, and does not require development of technical skills in stem cell culture. For a limited investment, both in terms of technique and time, this allows to relatively rapidly obtain lens organoids that recapitulate specific aspects of lens biology, with the added possibility of performing functional genetic analysis in a cost-effective manner. The 21EM15 organoids provides therefore the lens community with a compelling new model to improve the understanding of lens biology. Cataract remains a major public health problem that is currently only treated by surgery. It would therefore be interesting to develop drug treatments, particularly in order to offer alternatives to populations in countries that do not have ready access to surgery, or to prevent cataract formation in populations exposed to cataractogenic conditions. From a clinical point of view, these lens organoids should make it possible to develop screens for identifying compounds that impact the prevention and/or treatment of cataract.

## Methods Cell

### Culture

21EM15 (obtained from Dr. John Reddan, Oakland University, Michigan) and HaCat cells (ATCC) were cultured in DMEM with 4.5 g/L glucose, L-glutamine, and sodium pyruvate included (Life Technologies), 10% Fetal Bovine Serum (Eurobio), and 1% penicillin-streptomycin (Life Technologies). A253 cells (ATCC) were cultured in McCoy 5A with 4.5 g/L glucose, L-glutamine, and sodium pyruvate included (Life Technologies), 10% Fetal Bovine Serum (Eurobio), and 1% penicillin-streptomycin (Life Technologies). All three cell lines were cultured in 100 mm cell culture treated Petri dishes (Corning) with 10 mL of culture medium. The cells were grown at 37 °C in a water saturated atmosphere with 5% CO_2_. These cells grow well in these conditions and usually are 80% confluent after three days in culture (after 10% seeding). Cells were passaged three times per week.

### Organoid culture

Round-bottom 96-well plates were coated with poly(2-hydroxyethylmethacrylate) (Polyhema) (Sigma) 1 week before cell culture. Polyhema was first dissolved at 50 mg/mL in 95% EtOH. This stock solution was then diluted to 30 mg/mL with absolute ethanol. For coating, each well, except for the outermost ones, was filled with 50 μL of a 30 mg/mL solution of Polyhema before the plate was allowed to dry overnight. For culture, the outermost wells were filled with 200 μL PBS to avoid evaporation. The remaining wells were seeded with 10,000 cells and filled with 200 μL culture medium for 10 days prior to experiments.

### Histology and image acquisition

For histological analyses, organoids were washed with 1X PBS, then fixed 24 hr in 4% pH 7 buffered formalin and processed for paraffin wax embedding in an Excelsior ES automaton (Thermo Scientific). Paraffin-embedded tissue was sectioned at 4 µm, mounted on positively charged slides and dried at 58°C for 60 minutes. Immunohistochemical staining was performed on the Discovery ULTRA Automated IHC stainer (ROCHE) using the Ventana detection kit (Ventana Medical Systems, Tucson, Ariz). For fluorescent labeling, following deparaffination with Discovery wash solution (Ventana) at 75°C for 8 minutes, antigen retrieval was performed using Ventana Tris-based buffer solution pH8 at 95°C to 100°C for 40 minutes. Endogen peroxidase was blocked with 3% H_2_O_2_ for 12 minutes. After rinsing, slides were incubated at 37°C for 60 minuteswith primary antibody. Signal enhancement was performed using a secondary HRP-conjugated antibody at 37°C for 16 minutes and DISCOVERY Rhodamine Kit (Roche) for 8 minutes. For fluorescence multiplex labelling, slides were prepared as follow. First Sequence: following deparaffination with Discovery wash solution (Ventana) at 75°C for 8 minutes, antigen retrieval was performed using Ventana proprietary, Tris-based buffer solution pH8, at 95°C to 100°C for 40 minutes. Endogen peroxidase was blocked with 3% H2O2 for 12 minutes. After rinsing, slides were incubated at 37°C for 60 minutes with primary antibody : rabbit anti-PROX1. Signal enhancement was performed using a Goat anti-Rabbit HRP at 37°C for 16 minutes and DISCOVERY Rhodamine Kit (542-568nm) for 8 minutes. Second Sequence: slides were neutralized with Discovery inhibitor (Ventana) for 8 minutes. After rinsing, slides were incubated at 37°C for 60 minutes with primary antibody mouse anti-JAG1. Signal enhancement was performed using a Goat anti-mouse HRP at 37°C for 16 minutes and DISCOVERY cy5 Kit for 8 minutes. Third Sequence: slides were neutralized with Discovery inhibitor (Ventana) for 8 minutes. After rinsing, slides were incubated at 37°C for 60 minutes with primary antibody Rabbit anti PAX6 Signal enhancement was performed using a Goat anti-rabbit HRP at 37°C for 16 minutes and DISCOVERY Fam Kit for 8 minutes. DAPI staining was used to visualize DNA/nucleus. For chromogenic labeling, following deparaffination with EZ Prep (Roche) at 75 °C for 8 minutes, antigen retrieval was performed using CC1 buffer (Roche) pH 8.0 at 95°C to 100°C for 40 minutes. Endogen peroxidase was blocked with 3% H_2_O_2_ for 12 minutes. After rinsing, slides were incubated at 37°C for 60 minuteswith primary Antibody. Signal enhancement was performed using a secondary HRP-conjugated antibody at 37°C for 16 minutes and revealed using the OmniMap DAB kit (Roche). The slides were counterstained with the Mayer’s hematoxylin. Antibodies were as follows: Ki67, ab16667 dilution 1/200 (Abcam); Cleaved Caspase 3, #9661 dilution 1/250 (Cell Signaling Technology); Lamin B1, A16909 dilution 1/200 (Abclonal); Laminin Z0097 dilution 1/200 (Dako); PAX6 AB2237 dilution 1/200 (Sigma-Aldrich); JAG1 sc390177 dilution 1/200 (Santa Cruz); PROX1 925202 dilution 1/200 (BioLegend); CRYAB ADI-SPA-223 dilution 1/200 (Enzo Life Science); HSF4 HPA048584, dilution 1/200 (Atlas Antibodies). TOMM20 ab186735 dilution 1/5000 (Abcam); Fibrillarin 32639 dilution 1/100 H2P2 (Cell Signaling Technology). HES staining was realized on a ST 5020 automaton (Leica). Bright field images were acquired using a digital slide scanner Nanozoomer (Hamamatsu), while fluorescence microscopy images were acquired using a DeltaVision Elite setup equipped with a Nikon IX71 microscope and a CoolSnap HQ camera (AppliedPrecision).

### Transparency, light focalization

Transparency was assessed by placing the organoids on an electron microscopy grid and imaging them with an AZ100 macroscope (Nikon). Light focalization was quantified using an Axio Observer inverted microscope (Zeiss). Briefly, a stack of images starting from the focus and progressively lowering the objective under the sample was acquired. For each image of the z-stack, the maximum light intensity at the center of the spheroid and the mean intensity in the field around the spheroid were quantified. The ratio of the maximum light intensity to the mean intensity was then calculated and plotted to give a graph.

### 21EM15 2D and 3D RNA isolation

RNAs were isolated using the Nucleospin kit (Macherey-Nagel) from a 100 mm Petri dish of 21EM15 cell culture at 80% confluence (2D) or 60 organoids (3D), for each replicate.

### Laser capture microdissection

For each replicate, 120 organoids were washed with 1X PBS and pelleted before being included in OCT and then snap frozen using a SnapFrost 80 deep freezer (Excilone). The frozen OCT block was then mounted onto a cryostat (Leica) and cut to obtain 10 µm sections. The sections were then deposited on <u>p</u>oly<u>e</u>thylene <u>n</u>aphthalate (PEN)membrane frame slides. OCT was removed by multiple washes: 2 washes in 70° Ethanol (-20°C; 5 min), 1 wash in 90% Ethanol (RT; 20 min), 1 wash in 100% Ethanol (RT; 20 min) and 3 washes in 100 % Xylene (RT; 1 min). The internal or external regions of the spheroids were microdissected using a XT laser capture microdissection setup (Arcturus). The RNA was isolated from these samples using the Arcturus PicoPure kit (ThermoFisher).

### 3’-end RNA-sequencing (RNA-seq) and analysis

Libraries were prepared from the extracted RNAs using the QuantSeq 3’ mRNA-Seq library Kit (Lexogen). The 3’-end seq library were sequence (strand-specific, 150 bp) by the Illumina NovaSeq 6000. Quality of the sequence were validated by FastaQC, and only sense reads were used for the analysis. The RNA-seq data are available on the NCBI Gene Expression Omnibus (GEO) database under series GSE228547. The adaptor sequence and the poly(A) tail were trimmed from the raw sequences, and only read length superior to 20 nucleotides were retained. Trimmed sequences were aligned by the STAR software (STAR(v2.7.8a)) (Dobin et al., 2013) onto the mouse genome (GRCm38.p6). Only uniquely mapped reads were conserved for the analysis. Reads were associated to genes by FeatureCounts (v1.6.0) (Liao et al., 2014). For differential gene expression analysis, only genes with an expression >0.2 cpm (counts per million) were considered. The R package edgeR (v3.32.1) (Chen et al., 2016) was used to identify significantly differentially expressed genes (DEGs), with as cut-offs: |Fold Change| (FC) >2 (|logFC| > 1) and a False Discovery Rate (FDR) < 0.05.

### Gene expression data analysis

To determine the pattern of expression of the 2D or 3D DEGs in the mice embryonic lens, we used microarray data from the iSyTE 2.0 database (Kakrana et al., 2018) to identify genes that exhibit lens-enriched expression in normal lens development across stages E14.5 to P0. As described previously, comparison of global gene expression data between lens and whole embryonic body tissue (WB) allows estimation of lens-enriched expression. To compare the regional transcriptomic profile between organoid and lens we used previously generated RNA-seq data to identify DEGs with a expression profile specific to isolated FC or LEC (Zhao et al., 2019). These data correspond to WT mice at stage E14.5, E16.5, E18,5 and P0.5. The identification of genes with an expression profile specific to FC or LEC was based on cut-offs of P-value adjusted < 0.05 and |FC| > 2.

## Acknowledgements

We thank the Biosit platforms (Rennes, France) for access to their facilities, and notably MRic (microscopy) and H2P2 (histology). We particularly thank Alain Fautrel, Nicolas Mouchet (H2P2 facility) for fruitful discussions. L.P. was supported by a grant from Retina France. S.A.L. was supported by National Institutes of Health / National Eye Institute [R01 EY021505 and R01 EY029770].

## Legends to supplemental information

**Supplemental Table 1**

**Gene expression in 2D and 3D cultures, measured by 3’ end RNAseq.** Columns B-D, gene identification (Ensembl ID, Entrez ID, and gene symbol). Columns E, logFC, log2(expression in 2D cultures / expression in 3D cultures). logCPM, log 2 counts per million. Column G, F, F-test. Column H, PValue, p-value for the expression in 2D and 3D culture, Student’s t test. Column I, FDR, adjusted p-value, Benjamini-Hochberg adjustment. Column J, CatMap, a boolean for the gene being present in the CatMap database. Columns K-N, WT_2D_1 to WT_2D_4, gene expression values in 4 2D samples. Columns O-R, WT_3D_1 to WT_3D_4, gene expression values in 4 3D samples. WT_3D_1 to WT_3D_4, gene expression values in 4 3D samples. Columns S-T, logCPM_WT_2D and logCPM_WT_3D, log 2 counts per million in 2D and 3D samples. Column U, gene description.

**Supplemental Table 2**

**Gene expression in laser-microdissected external and internal regions.** Columns B-D, gene identification (Ensembl ID, Entrez ID, and gene symbol). Columns E, logFC, log2(expression in internal region / expression in external region). logCPM, log 2 counts per million. Column G, F, F-test. Column H, PValue, p-value for the expression in internal and external regions, Student’s t test. Column I, FDR, adjusted p-value, Benjamini-Hochberg adjustment. Column J, CatMap, a boolean for the gene being present in the CatMap database. Columns K-N, µdiss_WT_Inter_1 to µdiss_WT_Inter_4, gene expression values in 4 internal regions. Columns O-Q, µdiss_WT_Exter_1 to µdiss_WT_Exter_3, gene expression values in 3 external regions. Columns R-S, logCPM_µdiss_WT_Inter and logCPM_µdiss_WT_Exter, log 2 counts per million in internal and external regions. Column T, gene description.

**Supplemental figure 1.**
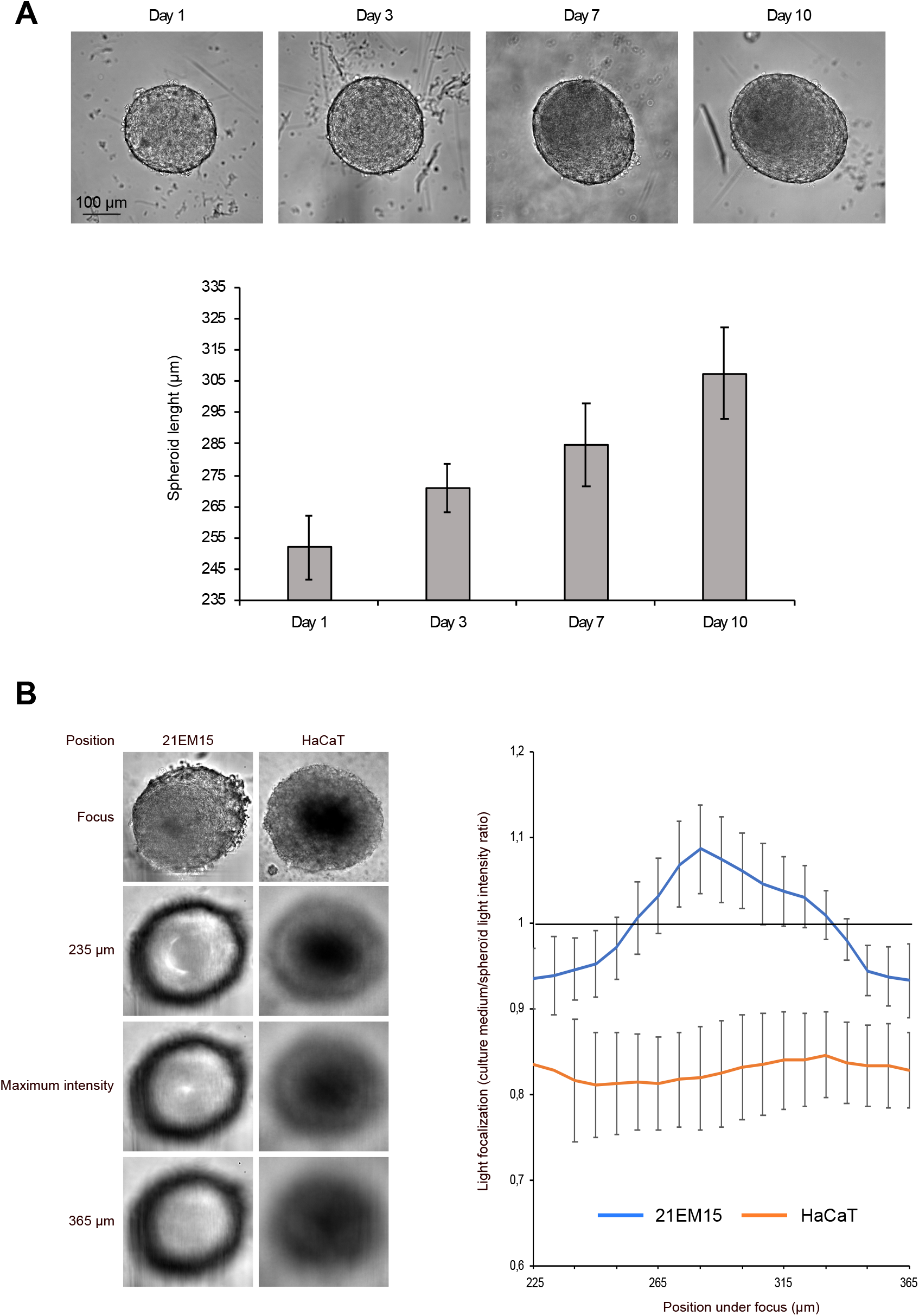
Growth pattern and light-focusing properties of 21EM15 spheroid. A, Wide field microscopy images and histogram showing the qualitative and quantitative analysis of the growth of 21EM15 spheroids over time. Graph is representative of three independent experiments with n=12 spheroids for each experiment. Error bars represent standard deviations. B, Microscopic images of 21EM15 and HaCat spheroids. Organoids derived from HaCaT cells are not able to focus light compared to organoids derived from 21EM15 cells. The graph on the right is representative of three independent experiments with n=12 spheroids for each experiment. Error bars represent standard deviations.

**Supplemental figure 2.**
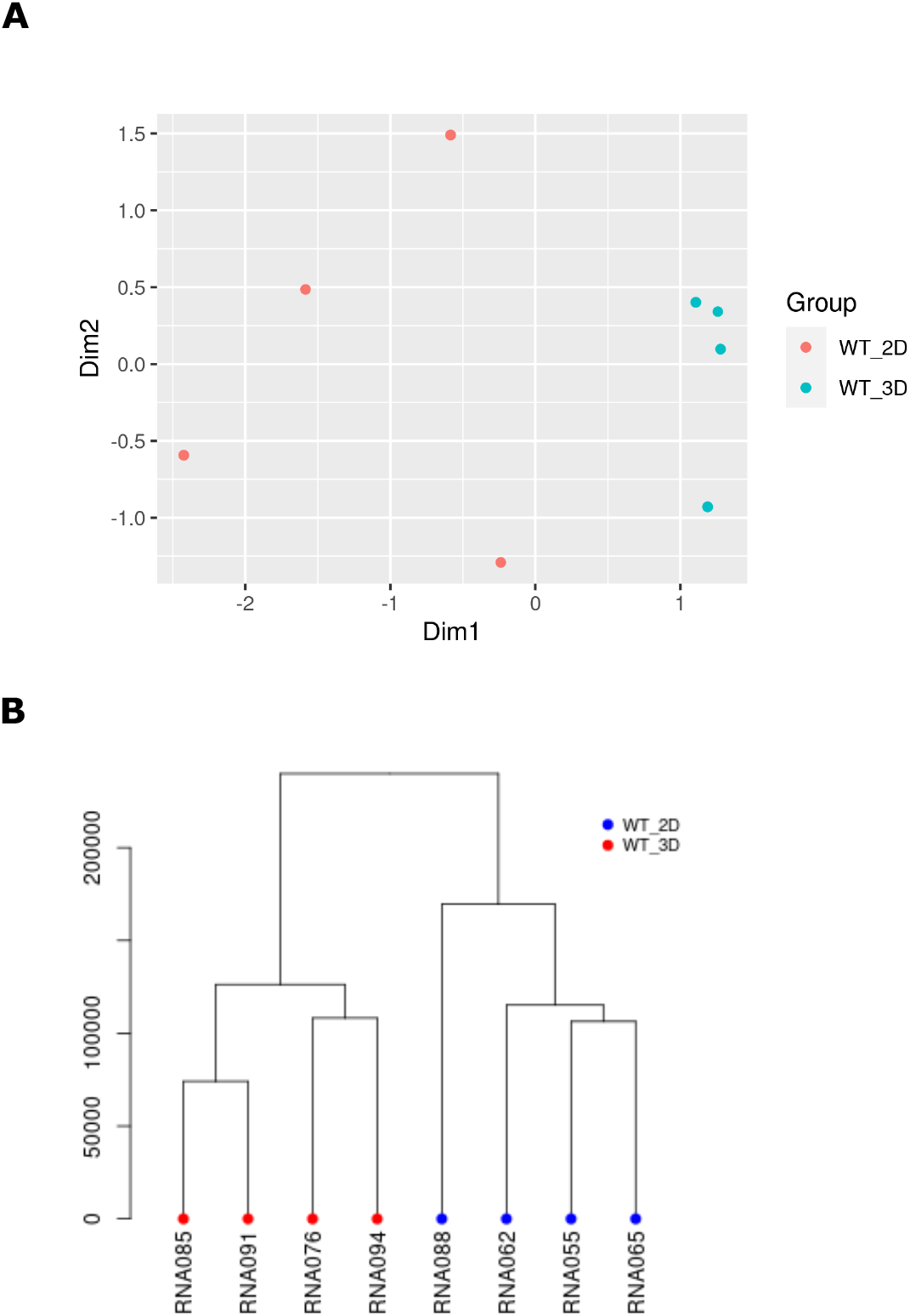
Quality control of RNA-seq of 21EM15 3D and 2D cultures. A, Principal Component Analysis (PCA) of RNA-sequencing (RNA-seq) datasets from lens culture. In the first dimension, the four samples from 3D cultures (blue) are grouped and separated from the four samples from 2D cultures (red). B, Dendrogram based on RNA-seq data. The four 3D samples (spheroids, red) and the 2D samples (blue) segregate into distinct clusters.

**Supplemental figure 3.**
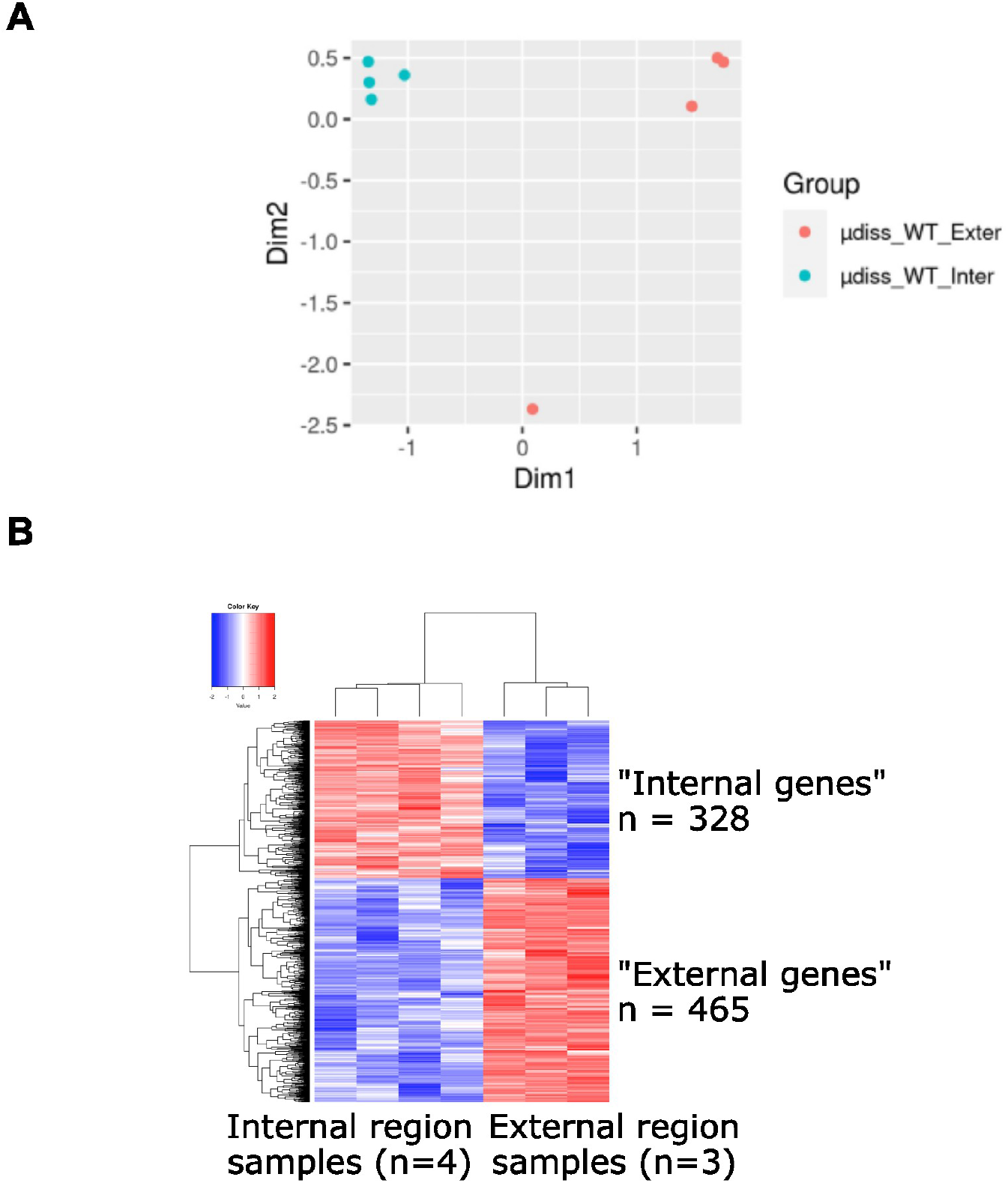
Quality control of RNA-seq of laser-captured microdissected 21EM15 spheroids. A, Principal Component Analysis of gene expression. The four samples microdissected from internal regions cluster together. Three out of four samples microdissected from external regions cluster together, and are clearly separated from the samples microdissected from internal regions in the first dimension. The fourth external region sample differs from the three other ones and we removed it for subsequent analyses. B, Heat map of the 793 DEGs, which separates “internal genes” from “external genes”.

